# Global Identification of Highly Expandable and Shrinkable Proteins via Intramolecular Distance Scoring Analysis

**DOI:** 10.1101/2025.03.21.643433

**Authors:** Tetsuji Okada, Fumiaki Tomoike, Takeshi Itabashi, Takayuki Nagae, Akira Nakamura

**Author notes:** Corresponding author: Tetsuji Okada, Department of Life Science, Gakushuin University, 1-5-1 Mejiro, Toshima-ku, Tokyo 171-8588, Japan.

## Abstract

Experimental and analytical characterizations of protein structural changes are essential for understanding various biological functions. It is well known that each protein or protein complex undergoes shape changes in a specific manner and to a certain extent, influenced by factors such as molecular interactions and the solvent environment. To enable quantitative comparisons of shape changes among multiple proteins, we previously introduced an index called the “UnMorphness Factor” (UMF), which is derived using a simple, original method— intramolecular distance scoring analysis (DSA). Using this approach, we have accumulated data for over 15,000 proteins. Here, we show that among the proteins exhibiting significant structural changes, some are found to undergo substantial isotropic volume expansion or contraction. The uniformity of three-dimensional size changes can be represented by a linear correlation between the average and standard deviation of all intramolecular C_α_-C_α_ distances across a protein’s structural set. Packing analysis of such proteins reveals that the void volume within the same polypeptide chain can increase by up to 50% relative to the reference state. Our study clearly demonstrates that this type of structural change plays a crucial role in understanding the detailed processes of ligand binding, catalytic reaction cycles, and protein folding pathways.

## Introduction

Proteins are structurally flexible and rely significantly on various non-covalent intramolecular interactions between the atoms of both the main chain and the side chains of the polypeptide. Their three-dimensional (3D) shapes are typically observed in the most energetically favorable state. However, the existence of multiple energetic minima can lead to different conformations for a given protein. Transitions between these conformations are often linked to the protein’s function in a physiological context. Some proteins and protein complexes undergo considerable structural changes, such as domain movement [1,2], domain swapping [3,4], and fold switching [5]. Even when the conformational change is not drastic, it can still have significant functional consequences.

Compared to these changes, isotropic volume changes are a type of structural change that is often overlooked. Although they are thought to influence various interactions both inside and outside the protein [6,7], a systematic and global analysis of this type of structural change has been lacking. Previous studies have shown that specific experimental conditions, such as temperature and structure analysis methods, affect the packing of polypeptides [8,9]. Additionally, the typical packing density of proteins and its variations have been examined using experimentally determined structures in the Protein Data Bank (PDB) [10,11]. However, these studies have not clarified the extent to which a specific protein undergoes isotropic volume changes under various experimental conditions.

We have been developing a method called Distance Scoring Analysis (DSA) for the reproducible quantitation of structural variability in proteins for which multiple experimental structures are available. By accumulating intramolecular C_α_-C_α_ distance data using DSA, we have illustrated the distribution of “morphable” proteins within an organism [12], and detected consistent *cis* bonds in each protein [13]. Naturally, isotropic volume changes in a protein, if they occur, are likely to be reflected in such intramolecular distance data. Here, we present the results of a global analysis of volume changes across more than 15,000 proteins, using approximately 160,000 chains from ∼120,000 PDB entries. Some highly expandable or shrinkable proteins identified in this study highlight the elasticity of naturally folded polypeptides from a volumetric perspective.

## Methods

### DSA analysis

DSA is a very simple yet unprecedented method for characterizing the structural variation of proteins for which at least three chains with PDB coordinates are available. For a given protein (single polypeptide), intramolecular distances between all pairs of C_α_-C_α_ atoms are calculated for each chain. Then, for each pair, the distances from multiple chains are used to compute a score by dividing the average distance by the standard deviation (stdev). The structural rigidity of a protein is indexed as the UnMorphable Factor (UMF), which is the average of all C_α_-C_α_ pair scores. Our previous study demonstrated that the UMF values for the majority of human and *E. coli* proteins fall within the range of 100–200 [12]. These calculations are performed using the original desktop Python tool, Score Analyzer [13] and a similar automated version implemented on the Google Colaboratory platform.

The main plot and the semi-log main plot of a protein represent the distribution of scores against the average distances. For proteins undergoing substantial structural changes, the semi-log main plot more clearly illustrates how distance variations extend to lower orders of scores. Another key plot from DSA is the progress plot, which depicts the change in the average score of a protein as the number of chains used increases. This plot can occasionally indicate which chain is structurally distinct from the others and is referenced to select pairs for comparisons of volume size and packing density.

As of March 15th, 2025, a total of 15,902 proteins have been analyzed using DSA, encompassing 164,992 chains from 124,109 PDB X-ray entries (the updated list is available at gses2.jp). This dataset excludes most proteins residing in large complexes and those with a significant number of missing C_α_ atoms in their full-length sequences. After obtaining intramolecular distance data for the dataset proteins, both the main plot and the semi-log main plot are archived. During the accumulation of these plot data, we occasionally identified proteins with very distinct patterns compared to the others, as described below. A closer examination of these proteins revealed that their data commonly exhibit signs of volume changes among the chains used in the analysis.

### Volume analysis

A variety of algorithms for protein volume calculation have been reported [14–21]. Regardless of the algorithm used, the comparison of two structural states of the same protein is unlikely to be affected, provided the same method is applied to both. In this study, we used the user-friendly online calculation tool ProteinVolume 1.3 [14], which provides three volume values (Total, Void, and Van der Waals (VDW)) as well as the packing density (VDW / Total) for the input polypeptide. Naturally, the VDW volume does not vary substantially as long as the same polypeptide chain, without any heteroatoms, is analyzed regardless of the structural state. Thus, differences in packing density between structural states can primarily be attributed to changes in void volume. In all cases presented in this study, the initial and final volume probe radii are 0.08 Å and 0.02 Å, respectively, with a minimum surface probe distance of 0.1 Å.

Structure superposition and graphics figure generation were performed using Discovery Studio Visualizer (BIOVIA). All plots were created using Igor Pro (WaveMetrics).

## Results and discussion

### Plots from DSA as Indicators of Volume Change

Our DSA results reveal intramolecular variations in C_α_-C_α_ distances for each protein (single polypeptide), represented by the main plot: score (average length / stdev) vs. average length. The main plot of a protein with non-specific structural variation often exhibits a round or ellipsoidal shape (Fig. 1 A, B), as seen in subtilisin Carlsberg from *Bacillus licheniformis* and ascorbate oxidase from *Cucurbita melopepo*. These proteins have been previously reported to exhibit high and low packing densities (0.783 and 0.708, respectively) [10]. Other examples of recent depositions are also presented in Supporting Figure 1.

**Fig. 1.**
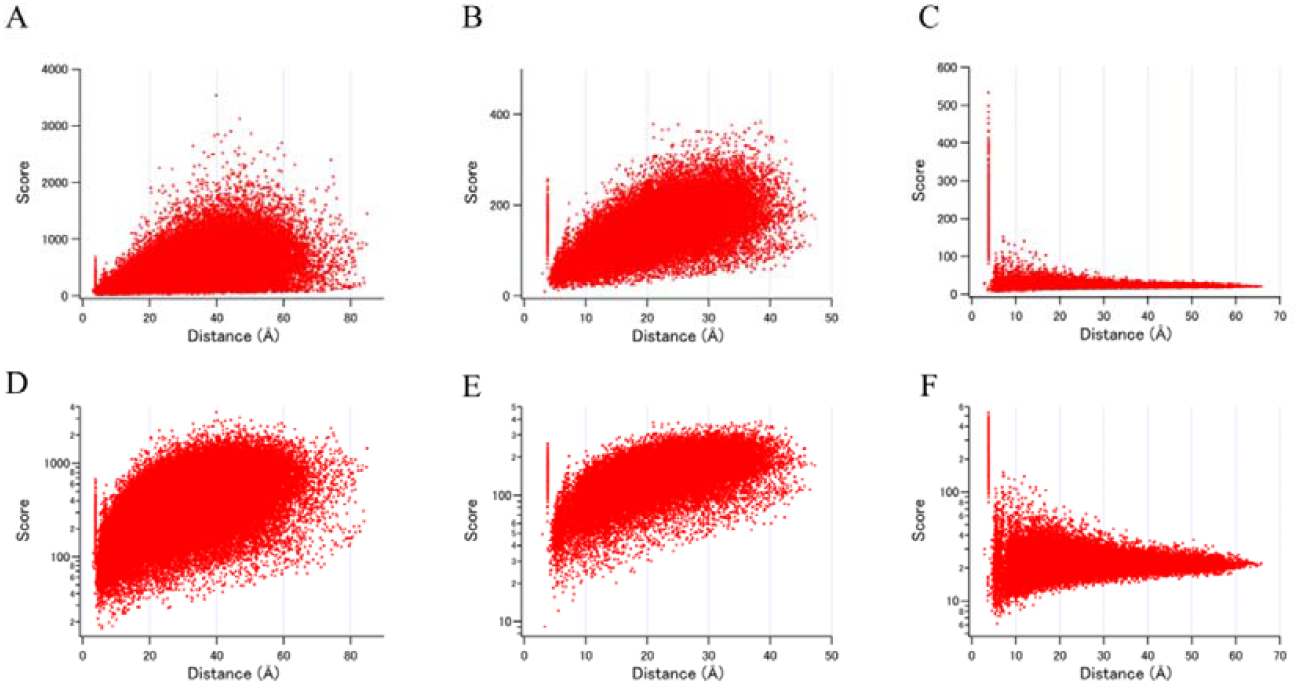
Comparison of the main plots (A ∼ C) and semi-log main plots (D ∼ F) of proteins with non-specific structural changes and those with isotropic volume changes. A, D: Subtilisin Carlsberg from *Bacillus licheniformis* (based on 25 chains from 25 entries) B, E: Ascorbate oxidase from *Cucurbita melopepo* (based on 8 chains from 4 enties) C, F: ChoX protein from *Rhizobium meliloti* (based on 10 chains from 5 enties)

Proteins with significant structural changes, such as domain motion, are better represented by the semi-log main plot, as demonstrated for calmodulin (Fig. 2). Conversely, some proteins exhibit distinct plot patterns, with a narrow score variation range in the long-distance region (Fig. 1C, F, and Fig. 3). This plot pattern is found to result from an almost linear correlation between average distances and standard deviations (Fig. 4). When the standard deviation is nearly proportional to the distance, the score value remains close to a constant, producing a sharp pattern in the semi-log main plot. To validate this interpretation, both visual (graphical) inspection and volume analysis were conducted on candidate proteins exhibiting significant volume changes. The list of these proteins is summarized in Table 1 and 2.

**Fig. 2.**
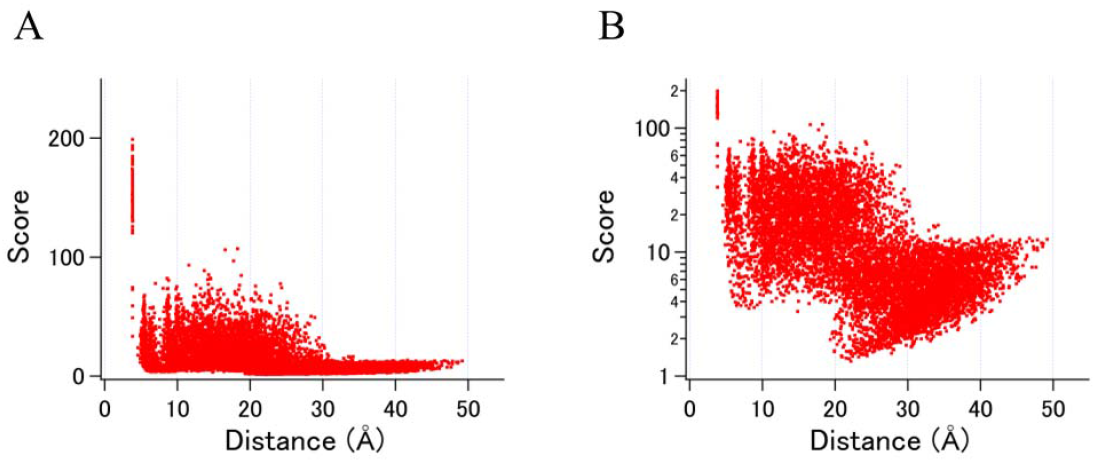
Examples of main plots for a protein with large domain movement. A. The main plot of human calmodulin. B. The semi-log main plot of human calmodulin. Both plots were generated using data from 89 chains from 89 entries.

**Fig. 3.**
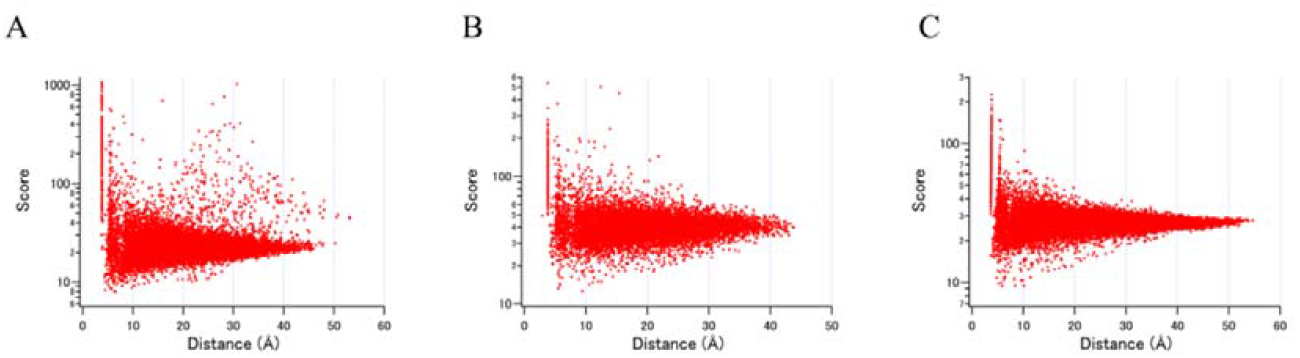
The semi-log main plots of proteins exhibiting volume changes. A. Pth protein from *Mycolicibacterium smegmatis* (based on 3 chains from 3 enties) B. BMEI0709 protein from *Brucella melitensis* (based on 5 chains from 2 enties) C. PD-L1 protein from *Phytolacca dioica* (based on 4 chains from 2 enties)

**Fig. 4.**
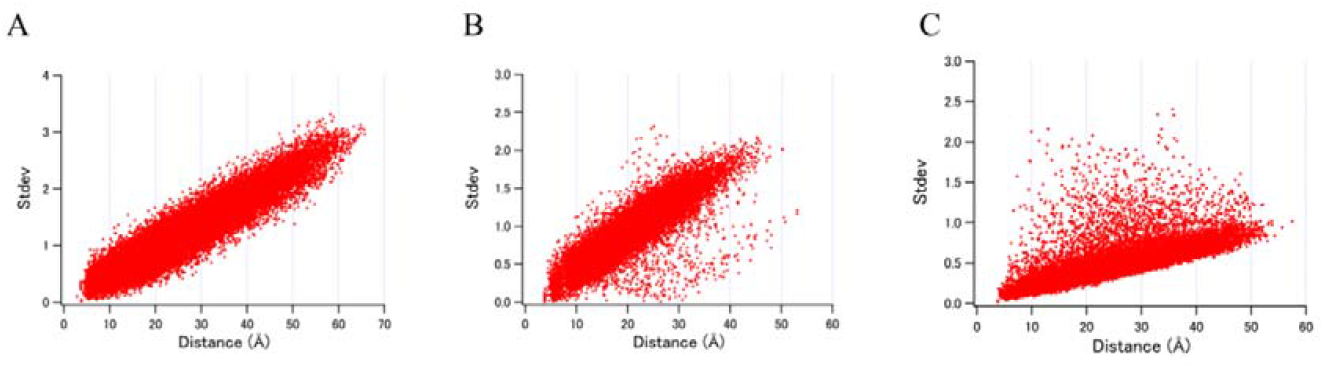
The stdev vs. average distance plots of proteins with volume changes. A. ChoX protein from *Rhizobium meliloti* (based on the same data as Fig. 1C) B. Pth protein from *Mycolicibacterium smegmatis* (based on the same data as Fig. 3A) C. Nsp16 protein from SARS-CoV-2 (based on 57 chains from 57 entries)

**Table 1.** Highly expanding proteins.

Highly expanding proteins
| Gene | Protein | UniProt | length | UMF | Pack. den. | Vol. ratio |
| --- | --- | --- | --- | --- | --- | --- |
| choX | Putative choline ABC transporter, periplasmic solute-binding comp. | Q92N37 | 287 | 24.1 | 0.654 | 1.516 |
| pth | Peptidyl-tRNA hydrolase | A0R3D3 | 191 | 27.8 | 0.662 | 1.509 |
| N.A. | 2'-O-methyltransferase | P0DTD1 | 296 | 57.1 | 0.667 | 1.461 |
| BMEI0709 | 4-hydroxyphenylacetate 3-monooxygenase | Q8YHT7 | 161 | 41.8 | 0.71 | 1.335 |
| cocE | Cocaine esterase | Q9L9D7 | 570 | 54.9 | 0.671 | 1.333 |
| mdlC | Benzoylformate decarboxylase (BFD) (BFDC) (EC 4.1.1.7) | P20906 | 523 | 107.3 | 0.684 | 1.292 |
| GAPDH | Glyceraldehyde-3-phosphate dehydrogenase | P00355 | 332 | 49.7 | 0.699 | 1.284 |
| map | Methionine aminopeptidase | Q9HXY1 | 258 | 41.1 | 0.699 | 1.281 |

Each length value represents the analyzed length, not the full length. The volume ratio (Vol. ratio) denotes the ratio of the void volume of the expanded chain to that of the reference chain. N.A. indicates "not applicable."

**Table 2.** Highly shrinking proteins.

Highly shrinking proteins
| Gene | Protein | UniProt | length | UMF | Pack. den. | Vol. ratio |
| --- | --- | --- | --- | --- | --- | --- |
| N.A. | Ribosome-inactivating protein PD-L1/PD-L2 | P84853 | 261 | 27.3 | 0.806 | 0.701 |
| CSNK2A2 | Casein kinase II subunit alpha' | P19784 | 324 | 90.2 | 0.777 | 0.780 |
| N.A. | Glutamine cyclotransferase | O81226 | 250 | 61.5 | 0.781 | 0.795 |
| eph | Epoxide hydrolase | Q1KLR5 | 236 | 76.9 | 0.767 | 0.830 |
| MA_3214 | Ulilysin | Q8TL28 | 259 | 83.7 | 0.774 | 0.831 |
| N.A. | plastic degrading hydrolase Ple629 | N.A. | 270 | 85.4 | 0.775 | 0.834 |
| ALB | Serum albumin (allergen Equ c 3) | P35747 | 580 | 57.7 | 0.77 | 0.850 |

Each length value represents the analyzed length, not the full length. The volume ratio (Vol. ratio) refers to the ratio of the void volume of the deviated chain to that of the reference chain. N.A. indicates "not applicable".

Following the volume analysis, it became evident that both expanding and shrinking proteins were identified, considering the average protein packing density of approximately 0.73∼0.74 [10,22]. We designate the pair of PDB chains as “reference” when their packing density is closer to the average, and as “expanded” or “shrunk” when their packing density is substantially lower or higher, respectively. Potential candidates for large volume changes are primarily identified based on the UMF value assigned to each protein. By definition, smaller UMF values indicate larger structural changes.

### Expanding proteins

In Table 1, we present a list of proteins for which at least one chain exhibits more than a 25% increase in void volume compared to the reference chain. For these proteins, the analyzed polypeptide length by DSA is more than 90% of the full length (Supporting Table 1). The average packing density of the reference chains for these eight proteins is 0.744, which is quite close to the reported average for many proteins [10,22]. Notably, the increase in void volume, calculated in the present work for the previously reported structure pairs from room-temperature and cryogenic-temperature crystallography [8] is mostly less than 10%, as shown in Supporting Table 2.

#### ChoX protein

The putative choline ABC transporter periplasmic solute-binding component from *Rhizobium meliloti* is identified as the most expandable polypeptide in our analysis, with a void volume increase of more than 50% (Table 1). As DSA uses the exact same range of polypeptide amino acid sequences from the experimentally available PDB coordinates of multiple chains for a given protein, we analyzed 90.3% of the full length of this protein (98.6% excluding the signal peptide region). Its remarkable degree of structural variation is well represented by the very low UMF value of 24.1 (the 13th lowest among 15,902 proteins, Supporting Table 3), though the structural change is not graphically obvious without domain movements. Instead, the change appears to be primarily an isotropic size change (Fig. 1F, 4A, 5A). Among the five PDB entries for this protein, two were assigned as semi-closed states and the other three as closed states [23,24]. Volume analysis of all ten chains (two chains per entry) is consistent with this assignment. Only the four chains from the semi-closed state (PDB IDs: 2REJ and 3HCQ, chains A and B) exhibit a void volume increase of 48.7– 51.6% compared to the reference chain (PDB ID: 2REG, chain A, with a packing density of 0.740). A 3D structural similarity search on the PDB site using the reference chain found 80 X-ray entries, including various extracellular substrate-binding proteins from other species, particularly members of the Pfam OpuAC family, to which ChoX also belongs. In contrast, only four entries of ChoX from *R. meliloti* were found for the expanded chain (chain A of 3HCQ), supporting the idea that the reference chain chosen here represents the normal packing state among other solute-binding proteins with a similar two-lobe fold.

**Fig. 5.**
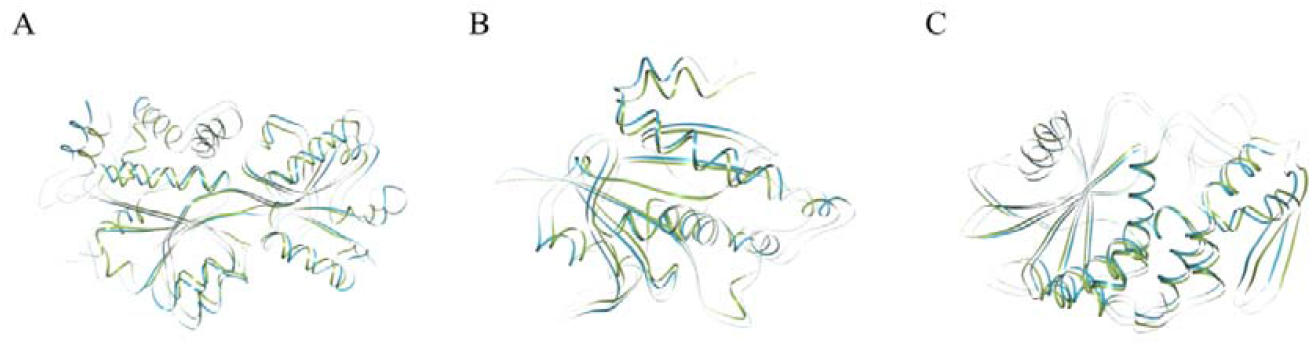
Superposed comparisons of expanded/shrunk chains with their respective references. A. ChoX protein from *Rhizobium meliloti*. Green: PDB ID 2REG, chain A; blue: PDB ID 3HCQ, chain B. B. Pth protein from *Mycolicibacterium smegmatis*. Green: PDB ID 3KJZ, chain A; blue: PDB ID 3KK0, chain A. C. PD-L1 protein from *Phytolacca dioica*. Green: PDB ID 3H5K, chain A; blue: PDB ID 3LE7, chain B. In all panels, the green structure represents the reference chain.

Previous structural studies on the lysine/arginine/ornithine-binding protein (LAO) from *Salmonella enterica*, a member of another two-lobe fold Pfam family (SBP_bac_3), have resulted in 23 PDB entries to date. As reported [25], our DSA analysis confirmed that LAO undergoes a bending-like conformational change upon ligand binding. In this case, the semi-log main plot pattern is typical for proteins undergoing domain movement (Supporting Fig. 2a), and the volume analysis of LAO does not show as remarkable a change as observed for ChoX.

At present, it is difficult to determine whether the unusual 3D expansion of ChoX is a unique feature, as no protein structures with more than 30% sequence identity to ChoX can be found in the current release of the PDB. The closest match is the glycine betaine-binding protein OpuAC from *Bacillus subtilis*, with a sequence identity of 28.28%. The domain movement of this protein is even less obvious than that of LAO (Supporting Fig. 2b).

#### Peptidyl-tRNA hydrolase (Pth)

The second most expanding protein identified in this study is Pth protein from *Mycolicibacterium smegmatis*, for which 100% of the full length was examined. There are only three X-ray chains available for this protein in the current release of the PDB (the minimum number required for DSA, which necessitates the calculation of standard deviation). In our experience, convergence of the UMF value for a given protein typically requires more than three chains. Thus, its unusually low UMF value of 27.8 is a rare case where nearly full structural variation has been explored with only three chains. The patterns of both the semi-log main plot and the standard deviation vs. average length plot are similar to those of the ChoX protein, indicating an isotropic shape change in 3D (Fig. 3A, 4B, and 5B).

The degree of void volume increase in the expanded chain (PDB ID: 3KK0) compared to the reference chain (PDB ID: 3KJZ) [26] is comparable to that found for ChoX. The remaining PDB entry (PDB ID: 3P2J) might also be considered as a reference, as its packing density (0.748) is close to that of 3KJZ (0.745). If we use this as the reference, the void volume increase in 3KK0 compared to 3P2J would be 1.529, which is higher than that of the ChoX protein. The three structures appear to have been obtained in the same crystal lattice but under different crystallization conditions: the precipitant composition of the expanded 3KK0 structure differs from that used for the other two structures. Thus, it is possible that the difference in solvent environment favored a partially folded state of this protein.

A close orthologue of this protein, Pth from *Mycobacterium tuberculosis* (82.2% identity for the full length), has also been structurally characterized, with eight PDB entries. However, we found no chains among them with a packing density of less than 0.74. Analysis of the physicochemical properties of this orthologue using ProtParam [27] reveals a significant difference in their instability indexes: the *M. smegmatis* protein is classified as stable, with an instability index of 28.76, while the

*M. tuberculosis* protein is classified as unstable, with an instability index of 44.48. Considering that the ChoX protein from *R. meliloti* also exhibits a low instability index of 23.78, the experimental detection of significant volume expansion in a protein might be more plausible if it has a somewhat stable nature, as indicated by the dipeptide composition used in the instability indexing algorithm [28]. It should also be noted, however, that two proteins (BMEI0709 and mdlC) from the top eight expanded proteins in Table 1 are not classified as stable according to the instability indexes from ProtParam.

The Pth protein from *M. smegmatis* was also characterized by solution NMR structure determination [29]. To investigate whether the structure ensemble shows any signs of volume change, we conducted DSA on the top three models from PDB entry 2NAF and found that the structural variation appeared to be randomly distributed across the entire chain (Supporting Fig. 3). Consistently, the packing density analysis of the models showed values ranging from 0.72 to 0.735. These results support the conclusion that the unusually expanded state of this protein is well beyond the fluctuation observable in the solution environment.

#### 2’-O-methyltransferase (nsp16)

Nsp16 from SARS-CoV-2 is the third most expanding protein identified in this study, with a void volume increase of approximately 45% in the product-bound conformation (PDB ID: 7LW3) and approximately 46% in the byproduct SAH-bound conformation (PDB ID: 7LW4), compared to the reference chain (PDB ID: 7L6T). While these structures were solved as complexes with the nsp10 protein, volume analysis was primarily conducted for the nsp16 component. Nsp10, which has an analyzed length of only 113 amino acids, also exhibits a substantial void volume increase of more than 25% in entries 7LW3 and 7LW4.

Most likely due to its biological importance in recent years, the number of PDB entries for this enzyme has increased to around 70, 57 of which have been analyzed by DSA. We began our analysis of this protein in August 2020 when only 17 entries were available. The addition of new data from subsequent entries resulted in a remarkable decrease in UMF, as demonstrated by the progress plot, and a substantial change in the pattern of the semi-log main plot (Fig. 6). This indicates that the two new chains (PDB IDs: 7LW3 and 7LW4) contributed to the change and likely differ from the previous ones in 3D size. This example clearly shows how we can identify new PDB chains whose coordinates contain information about the expansion or contraction of a protein, without the need to inspect each structure and its associated publication.

**Fig. 6.**
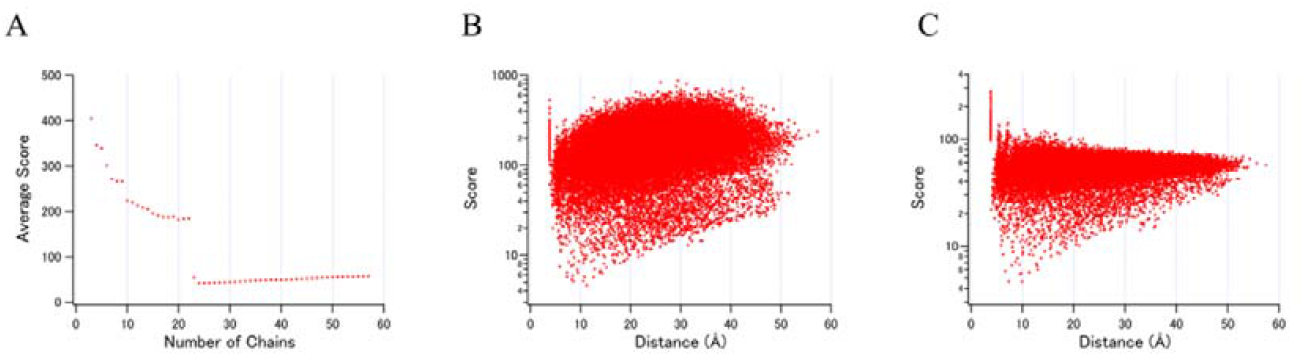
Plots from DSA obtained for the Nsp16 protein from SARS-CoV-2. A. The progress plot B. The semi-log main plot excluding data of expanded structures (based on 22 chains from 22 entries; up to entry 22 in A). C. The semi-log main plot including data of expanded structures (based on the same data as Fig. 4C).

In the case of nsp16, not only does isotropic size expansion occur, but it is also accompanied by other structural changes [30]. This is also reflected by the slight differences in the semi-log main plot and the standard deviation vs. average length plot, compared to those of ChoX and Pth proteins (Figs. 3, 4, and 6).

The fold pattern commonly found in the top three expanding proteins (ChoX, Pth, and nsp16) is the presence of interior β-strands surrounded by α-helices (Fig. 5). This feature is also consistent with some (but not all) of the other proteins listed in Table 1 and 2.

The abundance of deposited entries for nsp16 enabled us to examine the correlation between the resolution of the structures and the calculated packing density [31]. For the 57 chains used in the DSA, we observed nearly constant packing density values of approximately 0.74, regardless of the resolution. The only exceptions were two chains (PDB IDs: 7LW3 and 7LW4) (Supporting Fig. 4). This result supports the conclusion that the expanded form of nsp16 extends well beyond the variation observable among structures solved with differing electron density quality. Furthermore, it can be seen that the choice of reference chain does not significantly affect the conclusion regarding the large volume change observed in this study.

### Shrinking proteins

In Table 2, we present a list of proteins in which at least one chain exhibits a void volume decrease of 15% or more compared to the reference chain, with packing densities of approximately 0.77 or higher. For these proteins, the analyzed polypeptide length by DSA is more than 75% of the full length (Supporting Table 4). Overall, the degree of void volume change relative to the reference is less pronounced than that observed for the expanding proteins described above. This is likely because there is little removable void volume remaining within a folded polypeptide chain. In fact, it should be noted that the intramolecular atomic clash scores [32] appear to be worst for the structures with the highest packing density found for ligand-bound PD-L1.

#### Ribosome-inactivating protein PD-L1

Four chains from two PDB entries include the full-length PD-L1 protein from *Phytolacca dioica*. This protein exhibits isotropic 3D size shrinkage in its ligand-bound form (PDB ID: 3LE7) [33], with a void volume decrease of ∼30% in both chains compared to the reference structures (PDB ID: 3H5K) [34]. Its packing density of 0.806 is unusually high compared to the 0.748 observed in the reference chain. These two entries appear to have been obtained under nearly identical crystallization conditions and crystal lattice, yet with significantly different solvent contents. As observed with the three expanding proteins, both the semi-log main plot (Fig. 3C) and the stdev vs. average length plot of PD-L1 are consistent with isotropic structural size differences among the four chains.

DSA results are also available for homologous proteins with more than 80% sequence identity: antiviral protein S from *Phytolacca americana* and PD-L4 from *Phytolacca dioica*. Among these structures, the asymmetric unit of PD-L1 contains two polypeptides, while the asymmetric units of antiviral protein S and PD-L4 contain only one chain each. The packing density of apo PD-L1 is comparable to that of both the apo and adenine-bound forms of PD-L4 and antiviral protein S. However, adenine-bound PD-L1, in its dimeric asymmetric unit, exhibits an exceptionally compact structure. The sequence differences between PD-L1 and PD-L4 (81.61% identity) are widely distributed around the conserved ligand-binding core region (Supporting Fig. 5), suggesting an as-yet-unknown cooperative mechanism underlying the shrinkage observed in PD-L1.

#### Casein kinase II subunit alpha’ (CSNK2A2)

The human CSNK2A2 gene product protein has been studied in complexes with various small-molecule compounds, resulting in 32 PDB entries, of which 27 are used for DSA, covering 92.6% of the full length. The structure with a significant volume change is identified similarly to the method described above for the nsp16 protein: the progress plot clearly shows that the indenoindole-type inhibitor AR18-bound structure (PDB ID: 6HMD) [35] exhibits a substantially shrunken conformation.

Like nsp16, but more prominently, the uniform contraction in this protein is superimposed with other structural deviations, as clearly represented by the plots from DSA (Supporting Fig. 6). As a member of the large eukaryotic kinase (EK) family, various local structural rearrangements are known to occur in CSNK2A2 [36,37]. However, to the best of our knowledge, isotropic shrinking over the entire polypeptide has not been demonstrated for any proteins in the EK family.

It should also be noted that, while our analysis does not provide clear evidence of isotropic volume change in CSNK2A1, a close homologue of CSNK2A2, another EK family protein, human CDK2, shows a moderate degree of volume expansion, with an approximately 20% increase in void volume. Therefore, we propose that some other proteins with the EK-fold structure may also be candidates for exhibiting substantial volume change.

#### Glutamine cyclotransferase (QC)

The QC protein from *Carica papaya* (PQC) adopts a β-propeller structure with a pseudo five-fold axis. Two PDB entries (2FAW and 2IWA), reported independently [38,39], contain two and one chains, respectively. We analyzed 86.8% of the full length (93.6% of the chain excluding the signal peptide) of these chains.

While the patterns of the semi-log main plot and the stdev vs. average distance plot for this protein do not clearly indicate isotropic shrinkage as observed in PD-L1 (Supporting Fig. 7), volume analysis reveals a ∼20% smaller void volume in both chains from 2FAW compared to the reference chain in 2IWA. The packing densities of the 2FAW chains (0.781 and 0.780) are also substantially higher than the 0.743 observed in the reference chain.

No structures with more than 40% sequence identity to PQC are currently found in the PDB. The most similar protein, QC from *Zymomonas mobilis* (ZmQC) [40], with a sequence identity of 37.4%, was also analyzed using DSA. The three chains from two entries of ZmQC are nearly indistinguishable, resulting in a high UMF value of 338.7. Their packing densities fall within a narrow range of 0.737–0.740. Therefore, the comparison of packing densities between PQC and ZmQC indicates that the ∼0.78 packing density observed in the shrunk PQC chains is unusually high for a β-propeller structure with a pseudo five-fold axis.

## Conclusions

In the present study, based on the DSA of approximately 16,000 proteins, we demonstrated the first global analysis of protein volume changes. Some proteins were found to exhibit strikingly expanded or shrunken forms among the analyzed PDB chains (Fig. 5). Uniformity in 3D expansion or contraction is suggested by the linear correlation between the average and the standard deviation of all intramolecular C_α_-C_α_ distances in a set of structures for a given protein (Fig. 4). Void volume increases of up to 50% and decreases of up to 30% were observed, corresponding to packing densities as low as ∼0.65 and as high as ∼0.80, respectively, for proteins with amino acid lengths greater than 150 [22]. Whereas the previously reported packing density values may vary even for the same protein depending on calculation methods and conditions—such as the examined sequence range and the specific identity of PDB chains—the degree of volume change observed in this study likely exceeds previously known variations caused by fluctuations in the solution environment, differences between room-temperature and cryogenic crystallography, and structural determination via X-ray versus electron microscopy.

Many of these examples have emerged from PDB depositions over the past two decades following extensive studies on protein packing and calculation methods [10,11,15,16,19]. As the size and overall shape of highly volume-changing proteins identified here do not appear to be restricted to specific groups or families, it is plausible that further characterization of proteins with newly determined structures will reveal that isotropic size change is not an uncommon type of transition.

It should also be noted that we may have overlooked some proteins for which only two chains with substantial differences in 3D size are available, due to the current DSA method requiring at least three chains for calculations. Moreover, the recent remarkable advancements in single-particle cryo-electron microscopy may have already uncovered other examples of isotropic protein volume changes.

As previously hypothesized for some of the proteins described in this study, a uniformly expanded form of a protein could represent a distinctly closed conformation upon ligand binding (PDB IDs: 2REJ and 3HCQ), a folding intermediate (PDB ID: 3KK0), or a transient state during a catalytic reaction cycle (PDB IDs: 7LW3 and 7LW4). Future discoveries of additional examples will shed light on the common microscopic mechanisms underlying isotropic protein volume changes.

## Author Contributions

Tetsuji Okada, Fumiaki Tomoike, Takeshi Itabashi, Takayuki Nagae and Akira Nakamura designed research, performed research, contributed analytic tools, analyzed data, and wrote the paper.

