## Supplemental Figures for "Global Identification of Highly Expandable and Shrinkable Proteins via Intramolecular Distance Scoring Analysis"

Fig. 1:

Example main plots of proteins with non-specific structural changes.

(a) HL652_19860 protein from *Herbiconiux sp.* SALV-R1 obtained by DSA of 7 chains from 2 entries.
(b) ACTG1 protein from *Oryctolagus cuniculus* obtained by DSA of 4 chains from 3 entries.
(c) Tmel_1135 protein from *Thermosipho melanesiensis* obtained by DSA of 8 chains from 4 entries.

(a)	　　(b)	　　　(c)

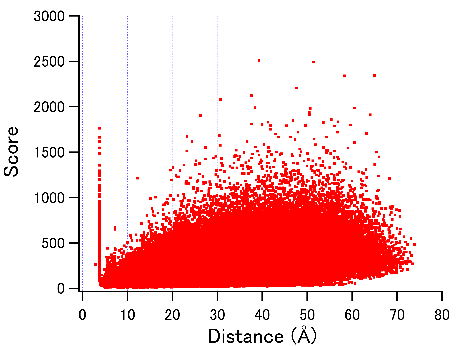

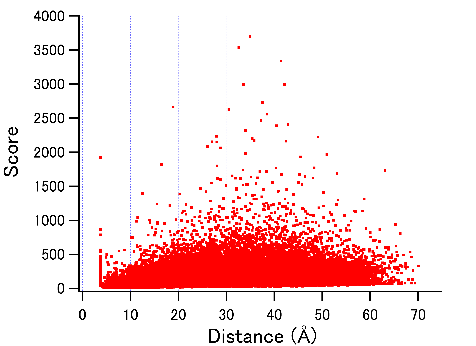

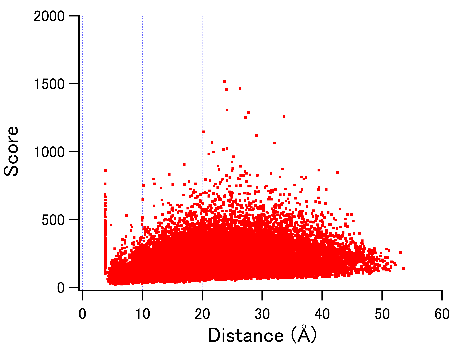

Fig. 2:

(a) The semi-log main plot of LAO protein from *Salmonella enterica* obtained by DSA of 22 chains from 22 entries.

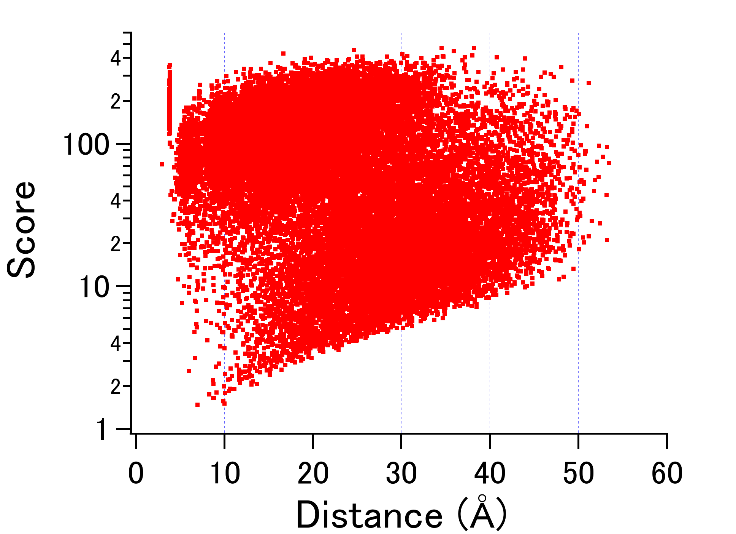

(b) The semi-log main plot of OpuAC protein from *Bacillus subtilis* obtained by DSA of 7 chains from 4 entries.

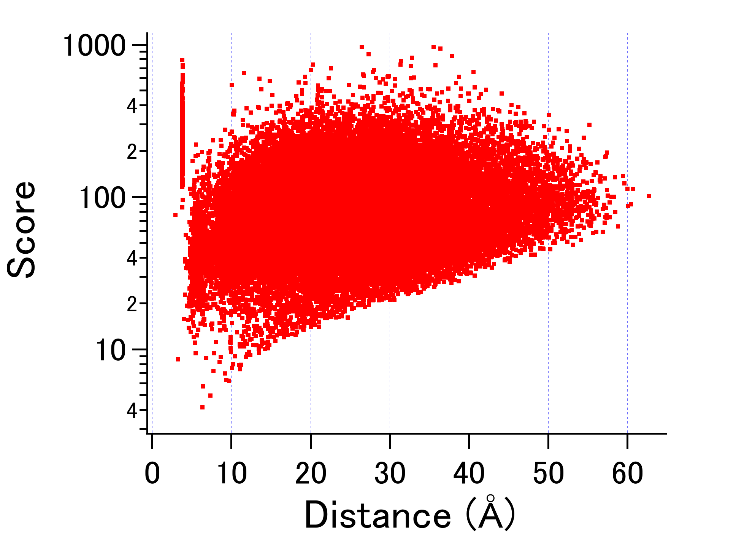

Fig. 3:

The semi-log main plot of pth protein from *Mycolicibacterium smegmatis* obtained by DSA of three NMR models of PDB entry 2NAF.

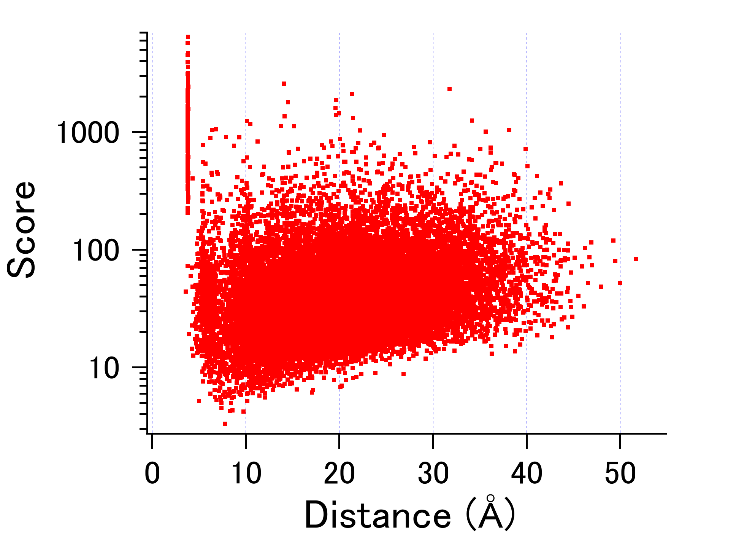

Fig. 4:

The correlation between packing density and resolution for 57 chains of nsp16. The two blue points represent the PDB IDs 7LW3 and 7LW4, while the green point represents the reference (7L6T).

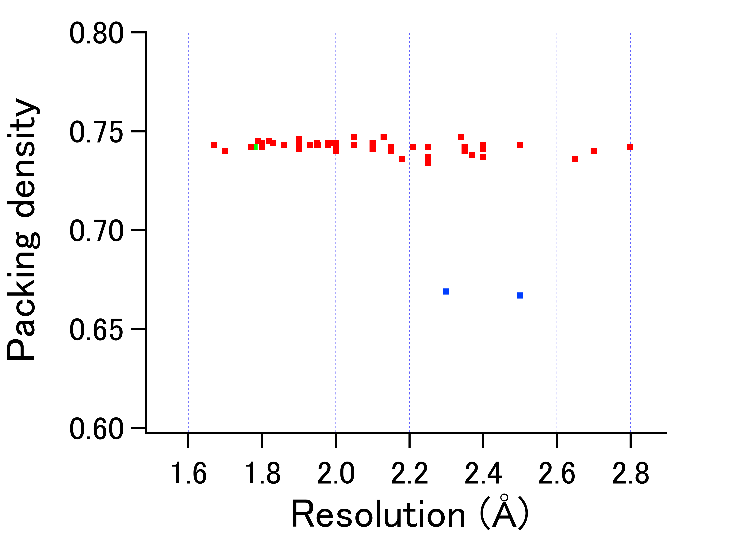

Fig. 5:

The sequence differences (shown in red) between PD-L1 and PD-L4 are marked on the reference chain (green) and the shrunken chain (blue) of PD-L1.

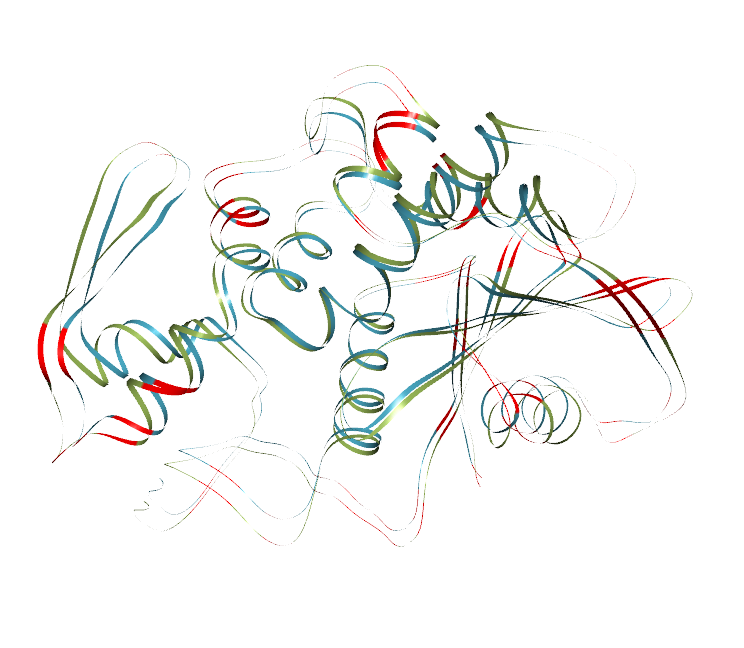

Fig. 6:

(a) The semi-log main plot and (b) the stdev vs. average distance plot of human CSNK2A2 protein obtained by DSA of 27 chains from 27 entries..

(a)

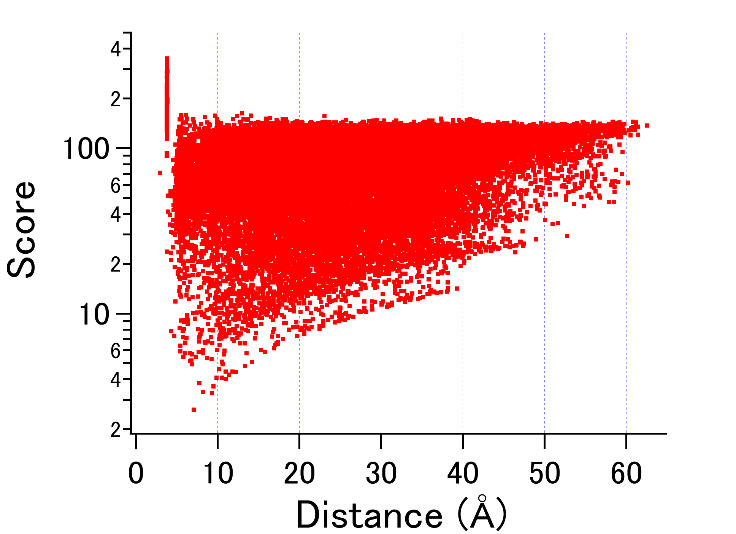

(b)

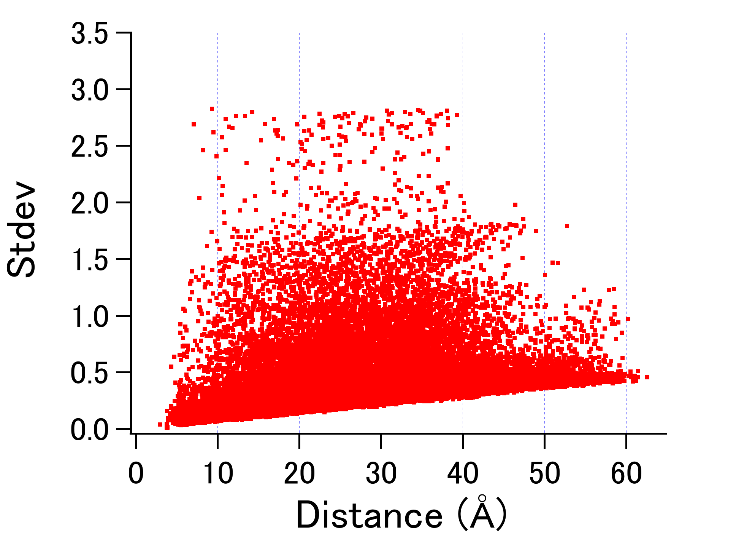

Fig. 7:

(a) The semi-log main plot and (b) the stdev vs. average distance plot of PQC protein from *Carica papaya* obtained by DSA of 3 chains from 2 entries.

(a)

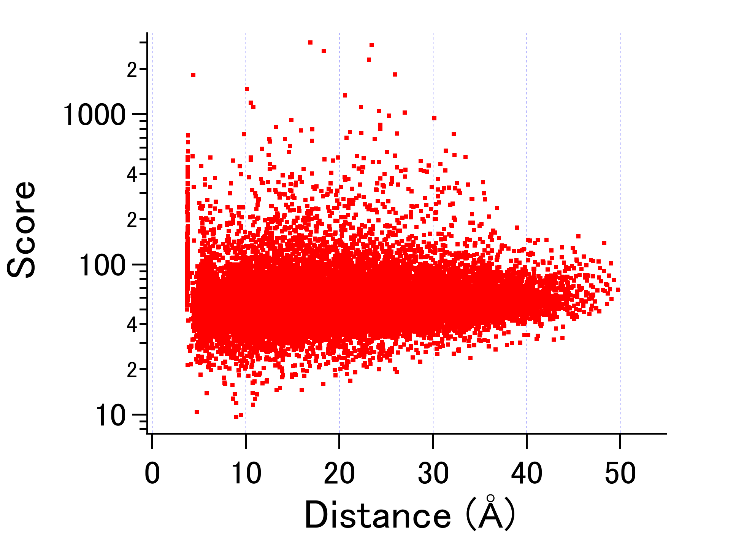

(b)

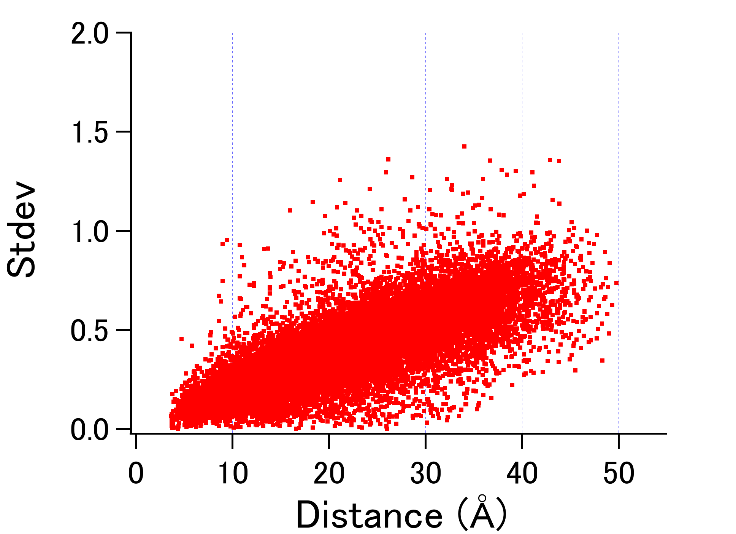

Supporting table 1

The details of the DSA results of the proteins in Table 1.

Supporting table 2

Volume and packing analysis of structure pairs determined at different temperatures.

Supporting table 3

The list of the 30 proteins with the lowest UMF values.

Supporting table 4

The details of the DSA results of the proteins in Table 2.
